# On automated model discovery and a universal material subroutine

**DOI:** 10.1101/2023.07.19.549749

**Authors:** Mathias Peirlinck, Kevin Linka, Juan A. Hurtado, Ellen Kuhl

## Abstract

Constitutive modeling is the cornerstone of computational and structural mechanics. In a finite element analysis, the constitutive model is encoded in the material subroutine, a function that maps local strains onto stresses. This function is called within every finite element, at each integration point, within every time step, at each Newton iteration. Today’s finite element packages offer large libraries of material subroutines to choose from. However, the scientific criteria for model selection remain highly subjective and prone to user bias. Here we fully automate the process of model selection, autonomously discover the best model and parameters from experimental data, encode all possible discoverable models into a single material subroutine, and seamlessly integrate this universal material subroutine into a finite element analysis. We prototype this strategy for tension, compression, and shear data from human brain tissue and perform a hyperelastic model discovery from twelve possible terms. These terms feature the first and second invariants, raised to the first and second powers, embedded in the identity, exponential, and logarithmic functions, generating 2^2×2×3^ = 4096 models in total. We demonstrate how to integrate these models into a single universal material subroutine that features the classical neo Hooke, Blatz Ko, Mooney Rivlin, Demiray, Gent, and Holzapfel models as special cases. Finite element simulations with our universal material subroutine show that it specializes well to these widely used models, generalizes well to newly discovered models, and agrees excellently with both experimental data and previous simulations. It also performs well within realistic finite element simulations and accurately predicts stress concentrations in the human brain for six different head impact scenarios. We anticipate that integrating automated model discovery into a universal material subroutine will generalize naturally to more complex anisotropic, compressible, and inelastic materials and to other nonlinear finite element platforms. Replacing dozens of individual material subroutines by a single universal material subroutine that is populated directly via automated model discovery—entirely without human interaction—makes finite element analyses more accessible, more robust, and less vulnerable to human error. This could forever change how we simulate materials and structures.

## 1 Motivation

Material modeling lies at the heart of a finite element analysis and selecting the appropriate material model is key to a successful finite element simulation [30]. A material model takes the local strains as input and calculates the stresses and their derivatives as output [27, 38]. Nonlinear finite element programs evaluate the material model locally, within every finite element, at each integration point, within every time step, at each Newton iteration [7, 39]. The local stresses and their derivatives then enter the global force vector and stiffness matrix to calculate the nodal displacements [30]. Finite element packages typically provide a comprehensive suite of built-in material models—-linear, polynomial, exponential, or logarithmic–with dozens of models to choose from [11, 23, 40, 41, 53]. This raises the question how we can select the best model and, probably more importantly, to which extent can we remove user bias throughout this selection process?

Admittedly, selecting the appropriate material model is a difficult task. This is especially true for unexperienced users or experienced scientists from other disciplines. For hyperelastic materials alone, commercial finite element packages offer the neo Hooke [60], Blatz Ko [6], Mooney Rivlin [42, 49], Yeoh [64], Gent [19], Demiray [12], Holzapfel [26], Ogden [45], and Valanis Landel [61] models, and continue to add new models as new releases emerge. To complicate matters, most finite element packages offer their users the flexibility to define their own custom-designed *user material subroutines* [1]. A user material subroutine is a modular software component that empowers the user to define and simulate complex material behaviors that cannot be captured by standard built-in material models. For example, our group has recently characterized different types of artificial meat and discovered material models that have never been used in any material library [55]. By implementing our own material subroutine, we can not only accurately model this complex material behavior, but also design and functionalize new materials [37, 43, 46]. This customization enhances the fidelity and accuracy of the simulation and enables the analysis of cutting-edge engineering problems for which standard material models fall short [25, 50, 55]. With this added flexibility in mind, do we now have to implement a new material subroutine every time we study a new material? And how do we discover the appropriate functional form that best describes the material behavior?

Recently, a trend has emerged to autonomously discover the model and parameters that best describe a specific material from experimental data, without any prior domain knowledge or user bias [5, 8]. There are several different strategies to achieve this. Most of them harness the power and robustness of algorithms developed for machine learning [3]. While some discover models that are interpretable, others do not.

*Non-interpretable approaches* closely follow traditional neural networks and typically discover functions of rectified linear unit, softplus, or hyperbolic tangent type [24]. The first representative of this category uses tensor basis Gaussian process regression, a special type of regression for isotropic hyperelastic materials that harnesses the representation theorem to a priori ensures objectivity [16]. In a rather abstract sense, it learns a 3 × 3 mapping that maps the three isotropic invariants onto the three coefficients of the stress tensor representation [17]. The second uses invariant-based constitutive artificial neural networks that a priori satisfy thermodynamic consistency by learning a free energy function from which they derive the stress [28, 33]. The third uses neural ordinary differential equations, special neural networks that a priori satisfy objectivity and polyconvexity by directly learning the derivatives of the free energy function that enter the stress definition [56, 58] While these approaches are straightforward, provide an excellent approximation of the data, and can be integrated manually within finite element software packages [24, 57], they learn non-interpretable models and parameters and teach us little about the underlying material.

*Interpretable approaches* discover models that are made up of a library of functional building blocks that resemble traditional constitutive models. The first representative of this category uses unsupervised learning and adopts sparse regression to discover interpretable models from a feature library of candidate functions [14, 15]. The second uses symbolic regression and genetic programming to discover mathematical expressions for invariant-based models in the form of rooted trees [2]. Here we combine all five approaches: We use a custom-designed invariant-based constitutive artificial neural network that a priori satisfies objectivity, thermodynamic consistency, and polyconvexity and autonomously discovers a free energy function that features popular constitutive terms and parameters with a clear physical interpretation [34–36]. In practice, interpretable models are limited by their functional form and approximate data less perfectly than non-interpretable models. At the same time, interpretable models are a generalization of popular existing constitutive models that–by design–translate seamlessly into user material subroutines.

The objective of this work is to integrate *automated model discovery* into the finite element workflow by creating a single *universal material subroutine* that automatically generates thousands of possible constitutive models. While we motivate this material subroutine from constitutive neural networks [35], the concept generalizes well to material models discovered via symbolic regression [2] or sparse regression or from feature libraries [14]. In Section 2, we briefly summarize the governing kinematic and constitutive equations. In Section 3, we introduce our constitutive neural network for automated model discovery. In Section 4, we translate all possible models of our network into a universal material subroutine and illustrate its pseudocode within the invariant-based `UANISOHYPER_INV` environment of the finite element package Abaqus. In Section 5, we illustrate the features of our user material subroutine by means of three types of examples: four benchmarks with popular constitutive models, two benchmarks with newly discovered models, and six realistic finite element simulations. We discuss our results in Section 6 and close with a brief conclusion and outlook in Section 7.

## 2 Governing equations

We begin by summarizing the governing kinematic and constitutive equations and reduce the general sets of equations to the special homogeneous deformations of uniaxial tension, uniaxial compression, and simple shear.

### 2.1 Kinematics

To characterize finite deformations, we introduce the deformation map ***φ*** that maps material particles ***X*** from the undeformed configuration to particles, ***x*** = ***φ***(***X***), in the deformed configuration [39]. We describe relative deformations within the sample using the deformation gradient ***F***, the gradient of the deformation map ***φ*** with respect to the undeformed coordinates ***X***, and its Jacobian *J*,

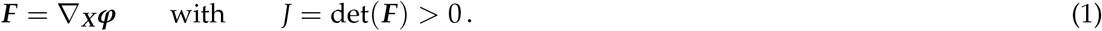

In the undeformed state, the deformation gradient is equal to the unit tensor, ***F*** = ***I***, and the Jacobian is equal to one, *J* = 1. A Jacobian smaller than one, 0 *< J <* 1, denotes compression and a Jacobian larger than one, 1 *< J*, denotes extension. To characterize an isotropic material, we introduce the three principal invariants *I*_1_, *I*_2_, *I*_3_ and their derivatives ∂_***F***_ *I*_1_, ∂_***F***_ *I*_2_, ∂_***F***_ *I*_3_,

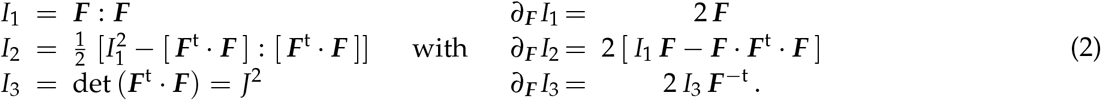

In the undeformed state, ***F*** = ***I***, the three invariants are equal to three and one, *I*_1_ = 3, *I*_2_ = 3, and *I*_3_ = 1. For isotropic, perfectly incompressible materials, the third invariant always remains identical to one, *I*_3_ = *J*^2^ = 1. This reduces the set of invariants to two, *I*_1_ and *I*_2_.

#### Tension and compression

For the case of uniaxial tension and compression, we stretch the specimen in one direction, *F*_11_ = *λ*_1_ = *λ*. For an isotropic, perfectly incompressible material with 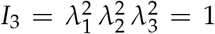, the stretches orthogonal to the loading direction are identical and equal to the square root of the stretch, *F*_22_ = *λ*_2_ = *λ*^−1/2^ and *F*_33_ = *λ*_3_ = *λ*^−1/2^. From the resulting deformation gradient, ***F*** = diag {*λ, λ*^−1/2^, *λ*^−1/2^}, we calculate the first and second invariants and their derivatives,

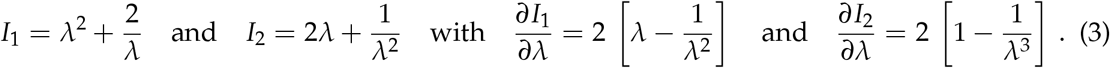

#### Shear

For the case of simple shear, we shear the specimen in one direction, *F*_12_ = *γ*. For an isotropic, perfectly incompressible material with *F*_11_ = *F*_22_ = *F*_33_ = 1, we calculate the first and second invariants and their derivatives,

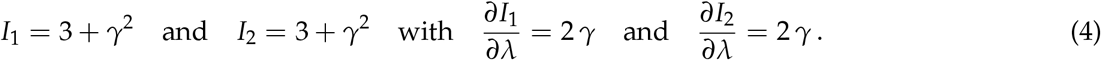

### 2.2 Constitutive equations

Constitutive equations relate a stress like the Piola or nominal stress ***P***, the force per undeformed area that is commonly measured in experiments, to a deformation measure like the deformation gradient ***F***. For a hyperelastic material that satisfies the second law of thermodynamics, we can express the Piola stress, ***P*** = ∂*ψ*(***F***)/∂***F***, as the derivative of the Helmholtz free energy function *ψ*(***F***) with respect to the deformation gradient ***F***, modified by a pressure term, − *p* ***F*** ^-t^, to ensure perfect incompressibility [39],

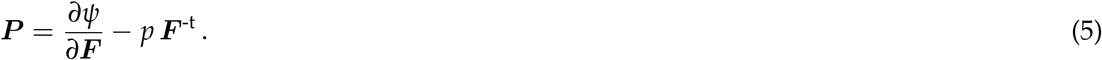

Here, the hydrostatic pressure, 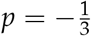 ***P*** : ***F***, acts as a Lagrange multiplier that that we determine from the boundary conditions. Instead of formulating the free energy function directly in terms of the deformation gradient *ψ*(***F***), we can express it in terms of the invariants, *ψ*(*I*_1_, *I*_2_), to yield the following expression for the Piola stress,

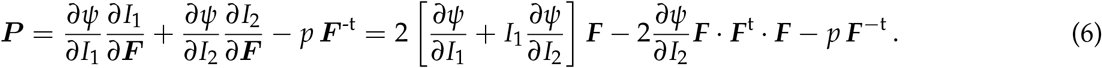

#### Tension and compression

For the case of uniaxial tension and compression, we evaluate the nominal uniaxial stress *P*_11_ using the general stress-stretch relationship for perfectly incompressible materials, *P*_*ii*_ = [∂*ψ*/∂*I*_1_] [∂*I*_1_/∂*λ*_*i*_] + [∂*ψ*/∂*I*_2_] [∂*I*_2_/∂*λ*_*i*_] − [1/*λ*_*i*_] *p*, for *i* = 1, 2, 3 with the invariants in tension and compression from equation (3). Here, *p* denotes the hydrostatic pressure that we determine from the zero stress condition in the transverse directions, *P*_22_ = 0 and *P*_33_ = 0, as *p* = [2/*λ*] ∂*ψ*/∂*I*_1_ + [2*λ* + 2/*λ*^2^] ∂*ψ*/∂*I*_2_. This results in the following explicit uniaxial stress-stretch relation,

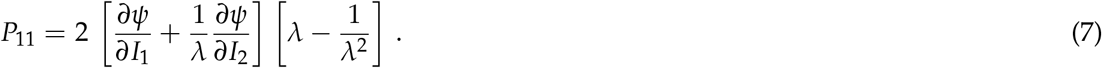

#### Shear

For the case of simple shear, we evaluate the nominal shear stress *P*_12_ using the general stress-stretch relationship for perfectly incompressible materials with the invariants for shear from equation (4). This results in the following explicit shear stress-strain relation,

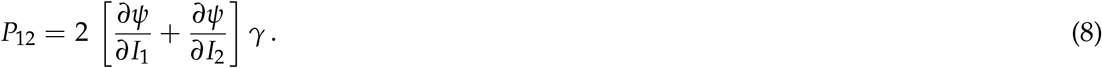

## 3 Neural network modeling

Motivated by these kinematic and constitutive considerations, we reverse-engineer a family of invariant-based neural networks that satisfy the conditions of thermodynamic consistency, material objectivity, material symmetry, incompressibility, constitutive restrictions, and polyconvexity by design [27, 39]. Yet, instead of building these constraints into the loss function [13, 48], we hard-wire them directly into our network input, output, architecture, and activation functions [34–36] to explicitly satisfy the fundamental laws of physics.

Figure 1 illustrates our constitutive neural network for isotropic, perfectly incompressible, hyperelastic materials. The network has two hidden layers with four and twelve nodes and a total of 24 weights. It takes the deformation gradient ***F*** as input and computes the first and second invariants [*I*_1_ − 3] and [*I*_2_ − 3]. The first layer applies activation functions *f*_1,1_ and *f*_1,2_ by gen-erating the first and second powers (∘)^1^ and (∘)^2^ of the two invariants, and multiplies them by the network weights *w*_1,1..12_. The second layer applies activation functions *f*_2,1_ and *f*_2,2_ and *f*_2,3_ by generating the identity (∘), the exponential function, (exp(∘) − 1), and the natural logarithm, (− ln(1 − (∘))), from these powers, and multiplies them by the network weights *w*_2,1..12_. The sum of all twelve terms defines the strain energy function *ψ*(***F***), from which the network calculates its output, the Piola stress, ***P*** = ∂*ψ*/∂***F***. Importantly, the networks is only selectively connected to a priori satisfy the condition of polyconvexity. The set of equations for this network takes the following explicit representation.

**Figure 1:**
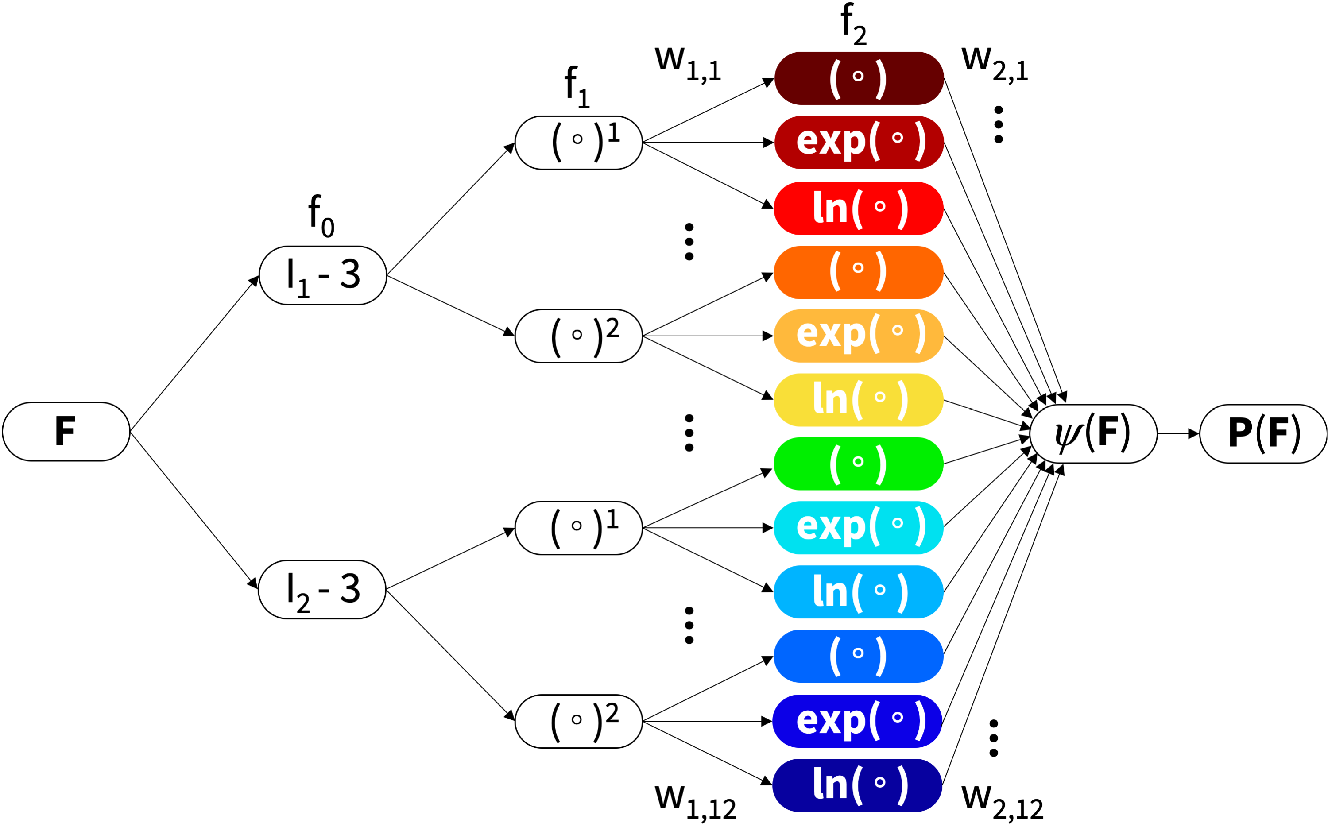
Constitutive neural network for isotropic, perfectly incompressible, hyperelastic materials. The network has two hidden layers with four and twelve nodes and 24 weights. It takes the deformation gradient ***F*** as input and computes the first and second invariants [*I*_1_ − 3] and [*I*_2_ − 3]. The first layer generates powers (∘)^1^ and (∘)^2^ of the two invariants and multiplies them by the network weights *w*_1,1..12_. The second layer applies the identity (∘), the exponential function, (exp(∘) − 1), and the natural logarithm, (− ln(1 − (∘))), to these powers, multiplies them by the network weights *w*_2,1..12_ and sums them up to calculate the strain energy function *ψ*(***F***), which defines the Piola stress, ***P*** = ∂*ψ*/∂***F***. The networks is selectively connected by design to a priori satisfy the condition of polyconvexity.

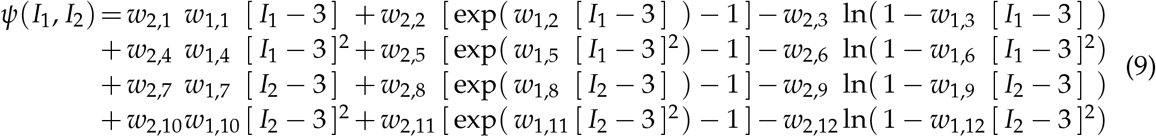

Here one of the first two weights of each row becomes redundant, and we can reduce the set of network parameters from 24 to 20, **w** = [(*w*_1,1_*w*_2,1_), *w*_1,2_, *w*_2,2_, *w*_1,3_, *w*_2,3_, (*w*_1,4_*w*_2,4_), *w*_1,5_, *w*_2,5_, *w*_1,6_, *w*_2,6_, (*w*_1,7_*w*_2,7_), *w*_1,8_, *w*_2,8_, *w*_1,9_, *w*_2,9_), (*w*_1,10_*w*_2,10_), *w*_1,11_, *w*_2,11_, *w*_1,12_, *w*_2,12_]. Using the second law of thermodynamics, we can derive an explicit expression for the Piola stress from equation (6), ***P*** = ∂*ψ*/∂*I*_1_ · ∂*I*_1_/∂***F*** + ∂*ψ*/∂*I*_2_ · ∂*I*_2_/∂***F*** − *p* ***F***^−t^,

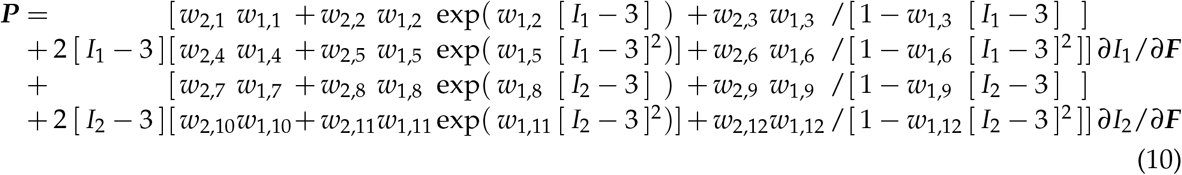

and correct it by the pressure term, − *p* ***F***^−t^, with 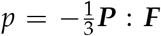. The constitutive neural network learns its weights, ***θ*** = {*w*_1..2,1..12_}, by minimizing a loss function *L* that penalizes the error between the model we want to discover and the experimental data. We characterize this error as the mean squared error, the *L*_2_-norm of the difference between model ***P***(***F***_*i*_) and data 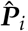, divided by the number of training points *n*_trn_,

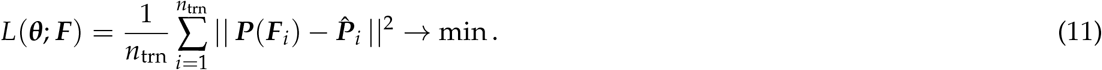

We train the network by minimizing the loss function (11) and learn the network weights ***θ*** = {*w*_1..2,1..12_} using the ADAM optimizer, a robust adaptive algorithm for gradient-based first-order optimization. To comply with physical constraints, we constrain all weights to always remain non-negative, *w*_*i,j*_ ≥ 0.

## 4 Universal material subroutine

Our objective is to create a seamless simulation pipeline from experiment–via discovered model and parameters–to simulation. To smoothly integrate our discovered model and parameters, we create a universal material subroutine that translates the local deformation, for example in the form of the deformation gradient ***F***, into the current stress, for example the Piola stress ***P***. This subroutine operates on the integration point level. Conveniently, some finite element codes provide the option to define a subroutine that works directly with the strain invariants and returns the first and second derivatives of the free energy function to calculate the stresses and their derivatives [1]. Towards this goal, we express the free energy function *ψ* from equation (9) in the following abstract form.

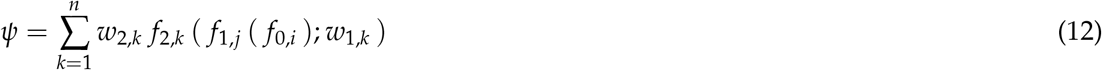

with

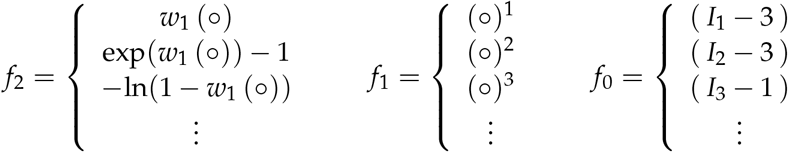

Here *f*_0_ maps the deformation gradient ***F*** onto a set of shifted invariants, [*I*_1_ − 3], [*I*_2_ − 3], [*I*_3_ − 1], [*I*_4_ − 1], [*I*_5_ − 1], that ensure that the strain energy function is zero in the underformed reference configuration, *f*_1_ raises these invariants to the first, second, or any higher order power, (∘)^1^, (∘)^2^, (∘)^3^, and *f*_2_ applies the identity (∘), exponential, (exp(∘) − 1), logarithm, − ln(1 − (∘)), or any other thermodynamically admissible function to these powers. The material subroutine can then calculate the Piola stress following equation (10) by using the derivatives of the individual functions *f*_0_, *f*_1_, *f*_2_.

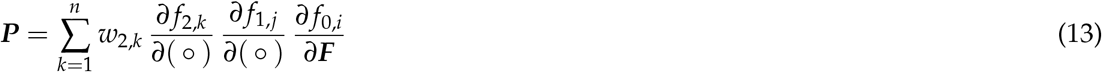

with

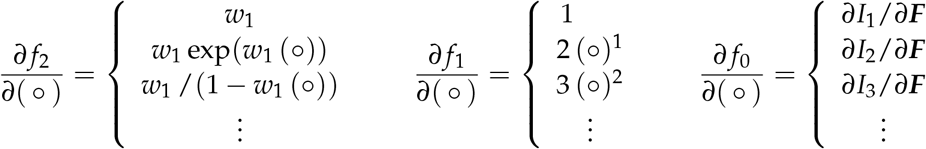

In implicit finite element codes that rely on a global Newton Raphson iteration, the material subroutine also calculates the second derivative for the tangent moduli.

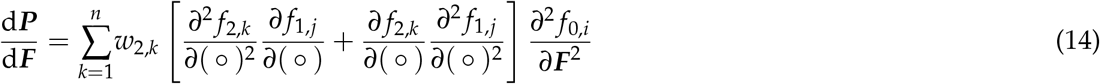

with

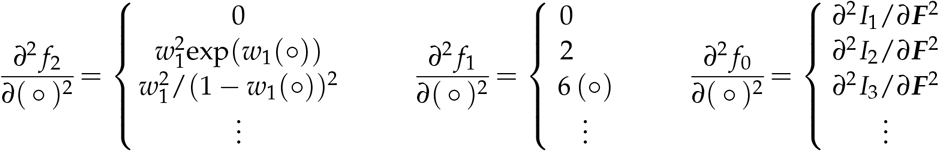

Figure 2 illustrates the free energy and stress contributions of our universal material subroutine for isotropic, perfectly incompressible, hyperelastic materials. The free energy function *ψ* is made up of twelve terms, based on the two invariants, [*I*_1_ − 3] and [*I*_2_ − 3], taken to the first and second powers, (∘)^1^ and (∘)^2^, and embedded in the identity (∘), the exponential function, (exp(∘) − 1), and the natural logarithm, (− ln(1 − (∘))). The odd rows illustrate these twelve terms, *ψ*(*I*_1_) and *ψ*(*I*_2_), for the special case of tension and compression with 0.5 ≤ *λ* ≤ 2.0, top four rows, and shear with 2.0 ≤ *γ* ≤ +2.0, bottom four rows. The even rows underneath each free energy term illustrate the corresponding Piola stress, ***P*** = ∂*ψ*/∂***F***, as the derivative of the free energy with respect to the deformation gradient, for the special case of tension and compression, top, and shear, bottom.

**Figure 2:**
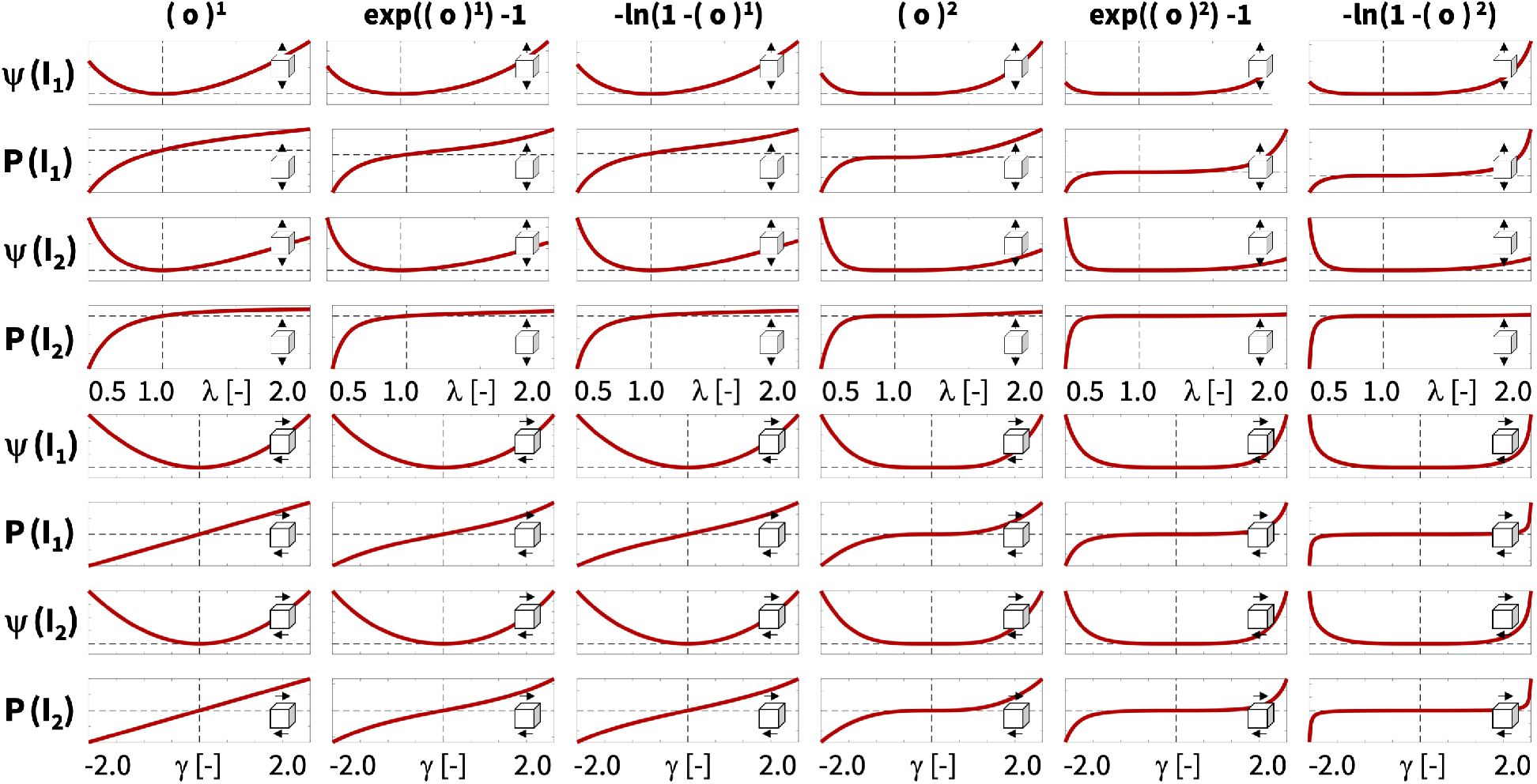
Universal material subroutine. Free energy and stress contributions. The free energy function *ψ* of our isotropic, perfectly incompressible, hyperelastic material is made up of powers (∘)^1^ and (∘)^2^ of the two invariants [*I*_1_ − 3] and [*I*_2_ − 3], embedded in the identity (∘), the exponential function, (exp(∘) − 1), and the natural logarithm, (− ln(1 − (∘))), odd rows. The stress ***P*** = ∂*ψ*/∂***F*** is the derivative of the free energy with respect to the deformation gradient, even rows. The twelve terms represent the twelve nodes of the constitutive neural network in Figure 1 for tension and compression with 0.5 ≤ *λ* ≤ 2.0, top four rows, and shear with −2.0 ≤ *γ* ≤ +2.0, bottom four rows.

Our discovered model translates seamlessly into a modular universal material subroutine within any finite element environment. Here we illustrate this translation by means of the Abaqus finite element analysis software suite [1]. We leverage the `UANISOHYPER_INV` user subroutine to introduce our discovered hyperelastic material strain energy function (9) or (12) in terms of the discovered pairs of network weights and activation functions. Specifically, our user subroutine defines the strain energy density, `UA(1)` = *ψ*, and the arrays of its first derivatives, `UI1(NINV)` = ∂*ψ*/∂*I*_*i*_, and second derivatives, `UI2(NINV`^∗^`(NINV+1)/2)`= ∂^2^*ψ*/∂*I*_*i*_∂*I*_*j*_, with respect to the invariants. With a view towards a potential generalization to anisotropic materials, we introduce an array of generalized invariants, `aInv(NINV)`= *I*∗_*i*_ with *i* = 1, …,`NINV`, where `NINV` is the total number of isotropic and anisotropic invariants, and adopt the `UANISOHYPER_INV` invariant numbering,

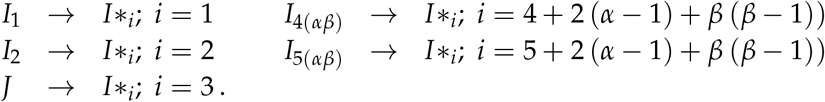

Here, *I*_4(*αβ*)_ = ***n*** _0*α*_ · (***F***^t^ · ***F***) · ***n*** _0*β*_ and *I*_5(*αβ*)_ = ***n*** _0*α*_ · (***F***^t^ ***F***)^2^ · ***n*** _0*β*_ are anisotropic invariants in terms of the unit vectors ***n*** _0*α*_ and ***n*** _0*β*_ that represent the directions of anisotropy in the reference configuration. For example, a transversely isotropic behavior with a single direction of anisotropy, *α* = *β* = 1, introduces two additional invariants, *I*_4_ = *I*_4(11)_ and *I*_5_ = *I*_5(11)_ [36].

Algorithm 1 presents the `UANISOHYPER_INV` pseudocode that describes how we compute the `UA(1)`, `UI1(NINV)`, and `UI2(NINV`^∗^`(NINV+1)/2)` arrays at the integration point level during a finite element analysis. In short, we begin by initializing all relevant arrays and read the activation functions *k f*_1,*k*_ and *k f*_2,*k*_ and weights *w*_1,*k*_ and *w*_2,*k*_ of the *n* color-coded nodes of our constitutive neural network in Figure 1 from our user-defined parameter table `UNIVERSAL_TAB`. Next, for each node, we evaluate its row in the parameter table `UNIVERSAL_TAB` and additively update the strain energy density function `UA`, its first derivative `UI1`, and its second derivative `UI2` using the node’s reference-corrected invariants `xInv`.

Algorithm 2 details the additive update of the free energy `UA` and its first and second derivatives

`UI1` and `UI2` within the user material subroutine `uCANN`.

Algorithms 3 and 4 provide the pseudocode for the two subroutines `uCANN_h1` and `uCANN_h2` that evaluate the first and second network layers for each network node with its discovered activation functions and weights.

### Algorithm 1

Pseudocode for universal material subroutine `UANISOHYPER_INV`

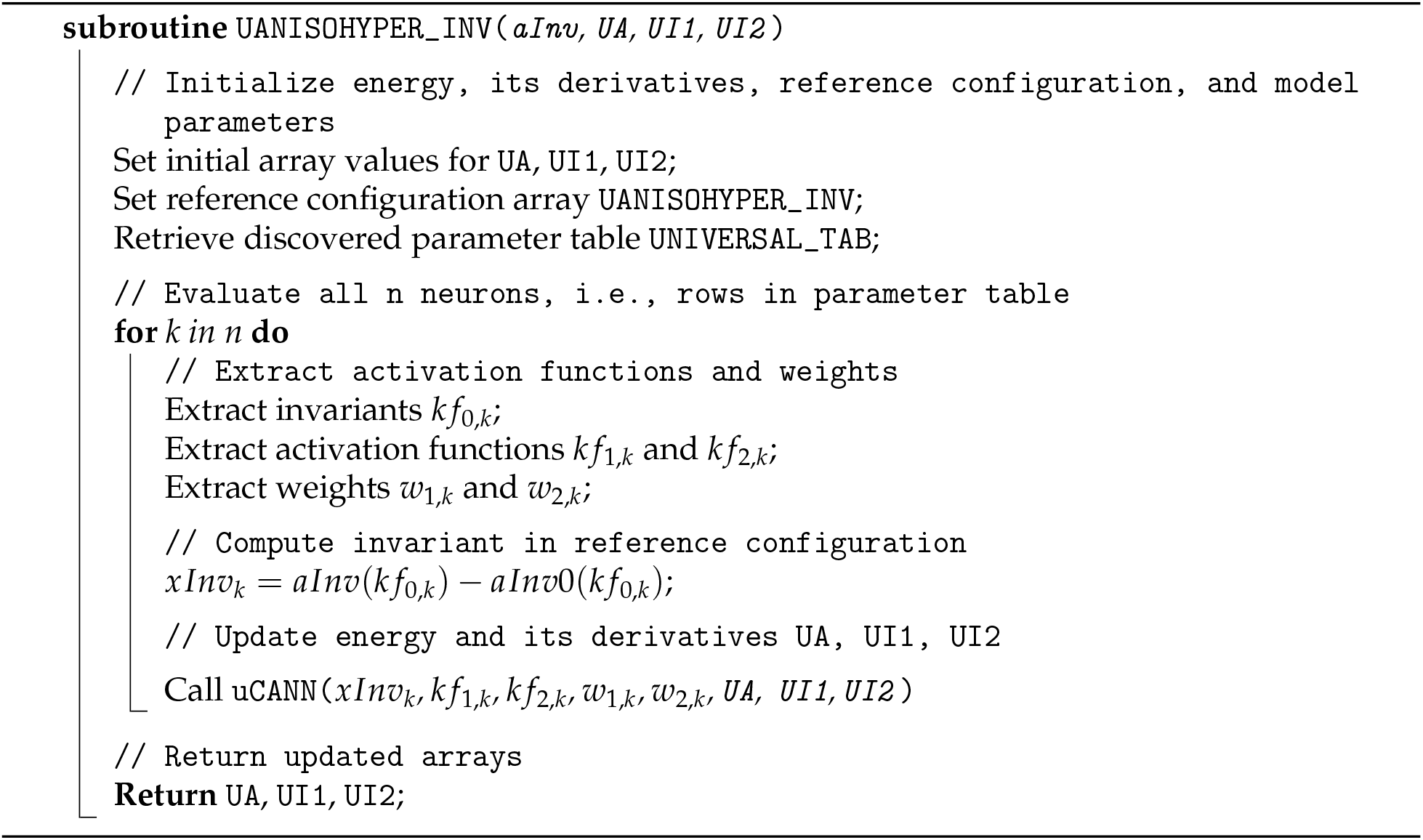

### Algorithm 2

Pseudocode to update energy and its derivatives for `UANISOHYPER_INV`

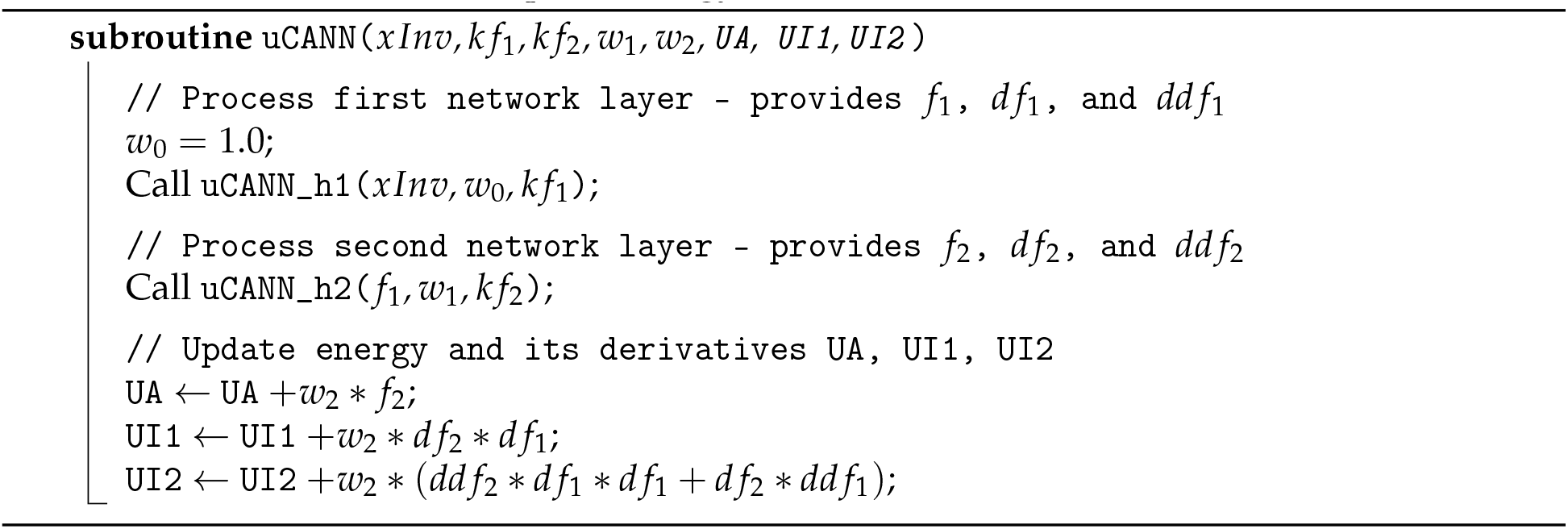

### Algorithm 3

Pseudocode to evaluate first network layer of `UANISOHYPER_INV`

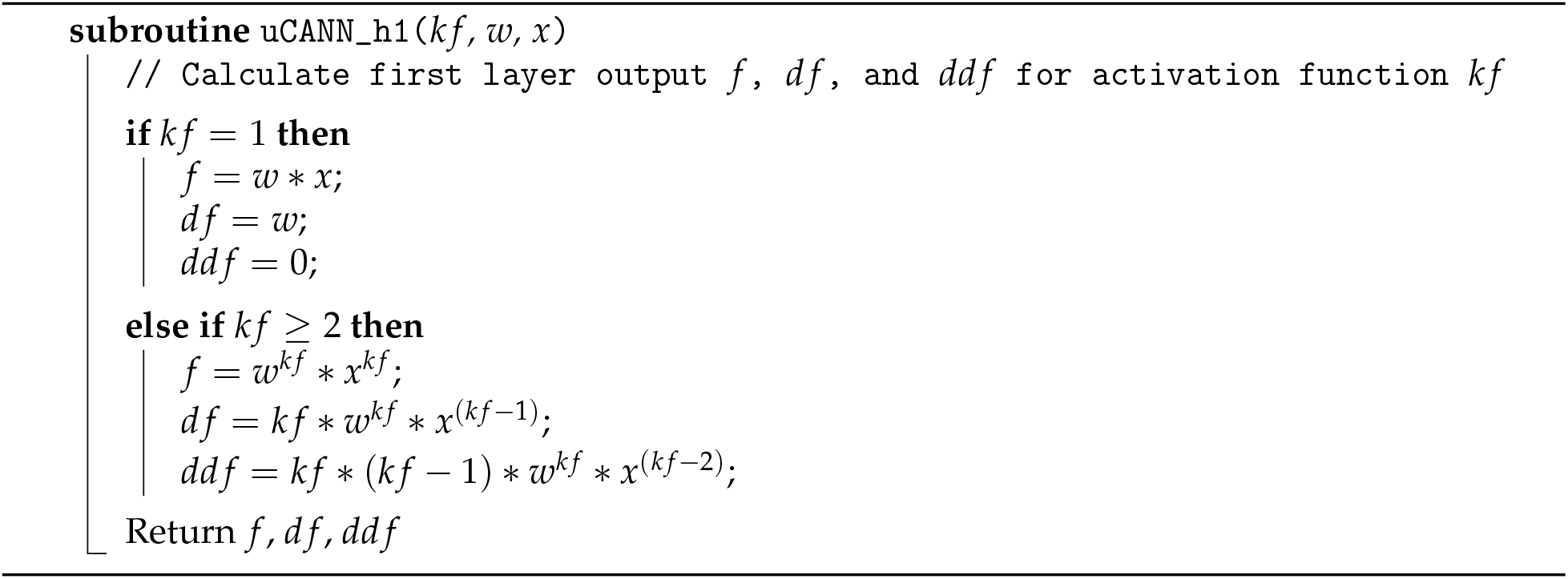

### Algorithm 4

Pseudocode to evaluate second network layer of `UANISOHYPER_INV`

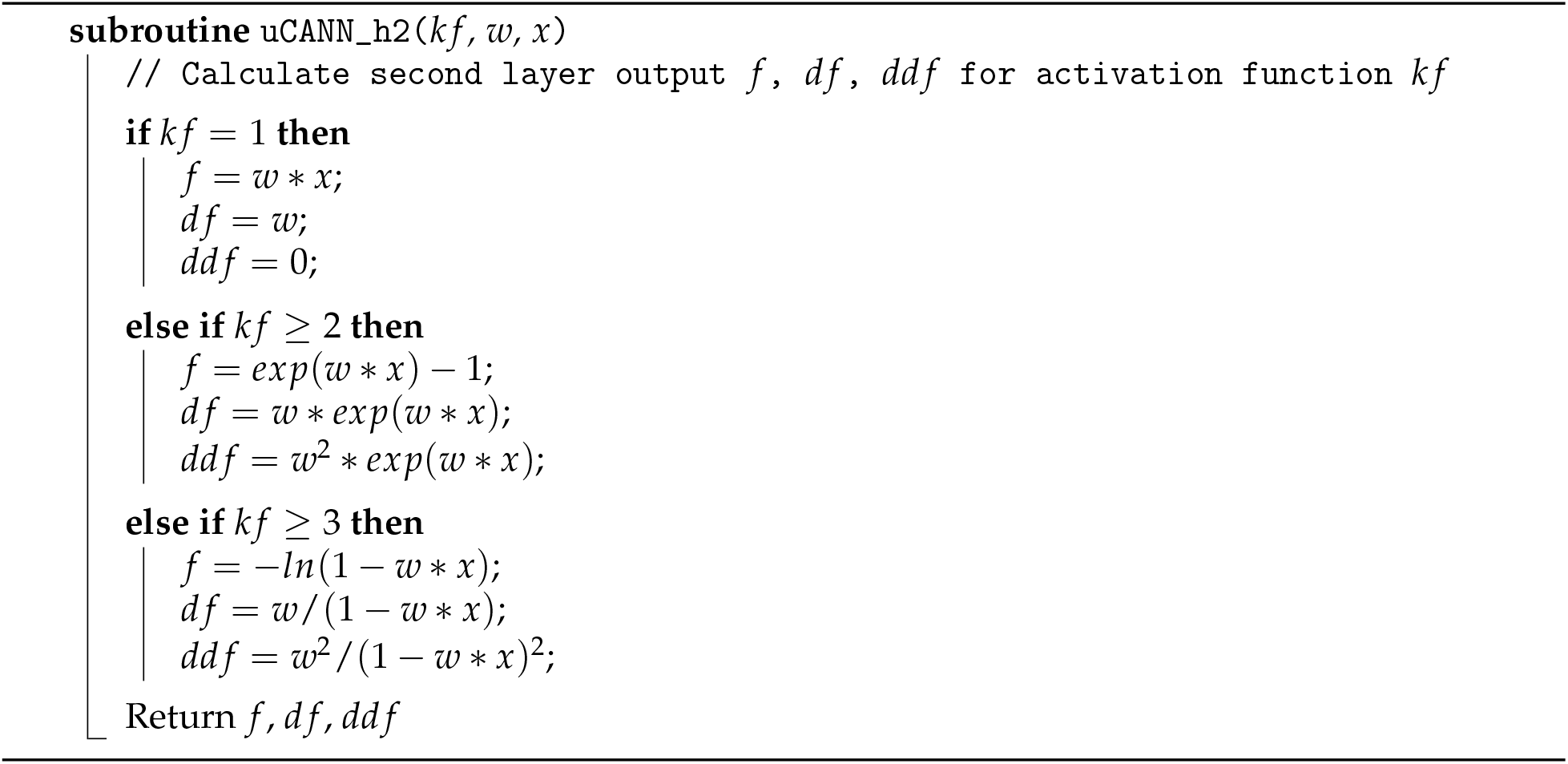

In Abaqus FEA, we define our model parameters in a parameter table; in our example, in the `UNIVERSAL_PARAM_TYPES.INC` file. Each row of this parameter table represents one of the colorcoded nodes in Figure 1 and consists of five terms, an integer *k f* 0 that defines the index of the pseudo-invariant *xInv*, two integers *k f* 1 and *k f* 2 that define the indices of the first- and second-layer activation functions, and two float values *w*1 and *w*2 that define the weights of the first and second layers.

> *PARAMETER TABLE TYPE, name=“UNIVERSAL_TAB”, parameters=5 INTEGER,, “Index of Pseudo-Invariant, kf0,o” INTEGER,, “Index of first hidden layer activation function, kf1,o” INTEGER,, “Index of second hidden layer activation function, kf2,o” FLOAT,, “Weight of first hidden layer, w1,o” FLOAT,, “Weight of second hidden layer, w2,o”

Within Abaqus FEA, we include the parameter table type definition using

> *INCLUDE, INPUT=UNIVERSAL_PARAM_TYPES.INC

at the beginning of the input file. We activate our user-defined material model through the command

> *ANISOTROPIC HYPERELASTIC, USER, FORMULATION=INVARIANT

followed by the discovered parameter table entries. For a fully activated constitutive neural network without any zero weights, the header and the twelve rows of this table reads as follows, where terms with zero weight can simply be excluded from the list.

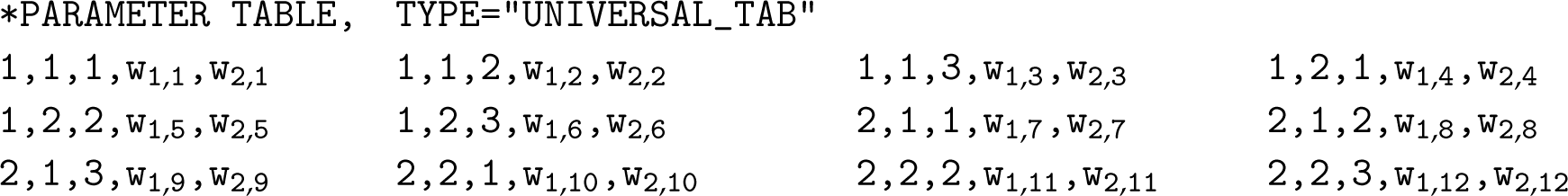

The first index selects between the first and second invariants, *I*_1_ or *I*_2_, the second index raises them to linear or quadratic powers, (∘)^1^ or (∘)^2^, and the third index selects between the identity, exponential, or logarithmic function, (∘), (exp(∘) − 1), or (− ln(1 − (∘))). Importantly, our user-defined material subroutine is universal by design. Combinations of the 2 × 12 weights naturally introduce popular and widely used material models as special cases. In the following, for illustrative purposes, we only highlight examples in terms of the first and second invariants *I*_1_ and *I*_2_. However, we can easily expand our user-defined material subroutine to include the third invariant *I*_3_ or any combination of the *I*_4*αβ*_ and *I*_5*αβ*_ invariants according to the invariant numbering scheme `NINV`. Moreover, the modular structure of our material subroutine facilitates a straightforward addition of additional first- and second-layer activation functions *k f* 1 and *k f* 2 within the `uCANN_h1` and `uCANN_h2` subroutines, or even completely novel layers with additional activation functions `uCANN_h*` within the hierarchical `uCANN` subroutine in Algorithm 2.

## 5 Results

We illustrate the features of our new user material subroutine in terms of three types of examples: First, we benchmark it with four popular constitutive models, demonstrate how to create the parameter tables for these models, and compare the simulations against the experimental data for gray matter tissue. Second, we benchmark it with two newly discovered models, create their parameter tables, and compare the simulations against both gray and white matter experiments. Finally, we demonstrate how it generalizes to realistic finite element simulations in terms of six different head impact simulations.

### 5.1 Benchmarking with popular constitutive models

To demonstrate that our universal material subroutine includes popular constitutive models as special cases, we benchmark our subroutine with four widely used models, translate their network weights *w*_1,•_ and *w*_2•,_ of the first and second layers into their model parameters, provide the material table for the input file to our material subroutine, and compare each simulation to experimental data. Table 1 summarizes our discovered non-zero gray matter weights *w*_1•,_ and *w*_2•,_ and model parameters *μ, μ*_1_, *μ*_2_, *a, b, α, β* when training our network with combined tension, compression, and shear data from human gray matter brain tissue [35].

**Table 1:** Neo Hooke, Blatz Ko, Mooney Rivlin, Demiray, Gent, and Holzapfel models and parameters. Discovered non-zero gray matter weights *w*_1•,_ and *w*_2•,_ and model parameters *μ, μ*_1_, *μ*_2_, *a, b, α, β* for training with combined tension, compression, and shear data from human gray matter brain tissue [35].

|  | neo Hooke<br>ten+com+shr |  | Blatz Ko<br>ten+com+shr |  | Mooney Rivlin<br>ten+com+shr |  | Demiray<br>ten+com+shr |  | Gent<br>ten+com+shr |  | Holzapfel<br>ten+com+shr |  |
| --- | --- | --- | --- | --- | --- | --- | --- | --- | --- | --- | --- | --- |
| | gray matter<br>$n = 15, 17, 35$ | | gray matter<br>$n = 15, 17, 35$ | | gray matter<br>$n = 15, 17, 35$ | | gray matter<br>$n = 15, 17, 35$ | | gray matter<br>$n = 15, 17, 35$ | | gray matter<br>$n = 15, 17, 35$ | |
| | $w_{1,\bullet}$<br>[-] | $w_{2,\bullet}$<br>[kPa] | $w_{1,\bullet}$<br>[-] | $w_{2,\bullet}$<br>[kPa] | $w_{1,\bullet}$<br>[-] | $w_{2,\bullet}$<br>[kPa] | $w_{1,\bullet}$<br>[-] | $w_{2,\bullet}$<br>[kPa] | $w_{1,\bullet}$<br>[-] | $w_{2,\bullet}$<br>[kPa] | $w_{1,\bullet}$<br>[-] | $w_{2,\bullet}$<br>[kPa] |
| $w_{\bullet,1}$ | 0.7880 | 1.1522 | — | — | 0.0026 | 0.4128 | — | — | — | — | — | — |
| $w_{\bullet,2}$ | — | — | — | — | — | — | 1.0529 | 0.8760 | — | — | — | — |
| $w_{\bullet,3}$ | — | — | — | — | — | — | — | — | 1.8399 | 0.4782 | — | — |
| $w_{\bullet,5}$ | — | — | — | — | — | — | — | — | — | — | 4.1833 | 4.7548 |
| $w_{\bullet,7}$ | — | — | 1.4156 | 0.6726 | 2.2122 | 0.4253 | — | — | — | — | — | — |
| | $\mu = 1.8159\text{kPa}$ | | $\mu = 1.9043\text{kPa}$ | | $\mu_1 = 0.0021\text{kPa}$<br>$\mu_2 = 1.8817\text{kPa}$ | | $a = 1.8447\text{kPa}$<br>$b = 1.0529$ | | $\alpha = 1.7597\text{kPa}$<br>$\beta = 1.8399$ | | $a = 39.7815\text{kPa}$<br>$b = 4.1833$ | |

#### Neo Hooke model

The neo Hooke model [60] is the simplest of all models. It has a free energy function that is constant in the first invariant, [*I*_1_ − 3], scaled by the shear modulus *μ*. We recover it as a special case from our network free energy (9) as

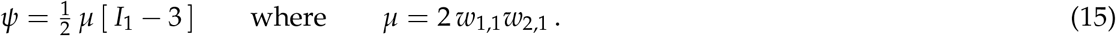

The neo Hooke model translates into the following material table for our universal material subroutine,

> *PARAMETER TABLE, TYPE=“UNIVERSAL_TAB” 1,1,1,w_1,1_,w_2,1_

and activates the first term of our model.

#### Blatz Ko model

The Blatz Ko model [6] has a free energy function that depends on the second and third invariants, [*I*_2_ − 3] and [*I*_3_ − 1], scaled by the shear modulus *μ* as 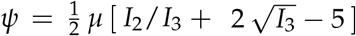. For perfectly incompressible materials, *I*_3_ = 1, we recover it as a special case of the network free energy (9) as

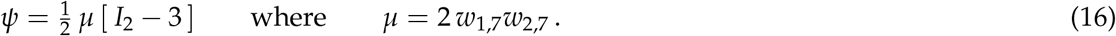

The Blatz Ko model translates into the following material table for our universal material subroutine,

> *PARAMETER TABLE, TYPE=“UNIVERSAL_TAB” 2,1,1,w_1,7_,w_2,7_

and activates the seventh term of our model.

#### Mooney Rivlin model

The Mooney Rivlin model [42, 49] is a combination of both free energy functions (15) and (16). It accounts for the first and second invariants, [*I*_1_ − 3] and [*I*_2_ − 3], scaled by the moduli *μ*_1_ and *μ*_2_ that sum up to the overall shear modulus, *μ* = *μ*_1_ + *μ*_2_. We recover it as a special case of the network free energy (9) as

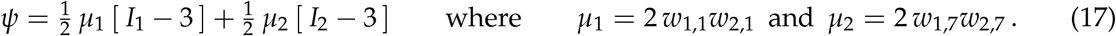

The Mooney Rivlin model translates into the following material table for our universal material subroutine,

> *PARAMETER TABLE, TYPE=“UNIVERSAL_TAB” 1,1,1,w_1,1_,w_2,1_ 2,1,1,w_1,7_,w_2,7_

and activates the first and seventh terms of our model.

#### Demiray model

The Demiray model [12] uses linear exponentials of the first invariant, [*I*_1_ − 3], in terms of two parameters *a* and *b*. We recover it as a special case of the network free energy (9) as

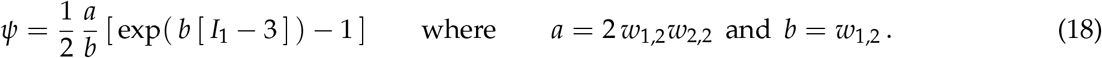

The Demiray model translates into the following material table for our universal material subroutine,

> *PARAMETER TABLE, TYPE=“UNIVERSAL_TAB” 1,1,2,w_1,2_,w_2,2_

and activates the second term of our model.

#### Gent model

The Gent model [19] uses linear logarithms of the first invariant, [*I*_1_ − 3], in terms of two parameters *α* and *β*. We recover it as a special case of the network free energy (9) as

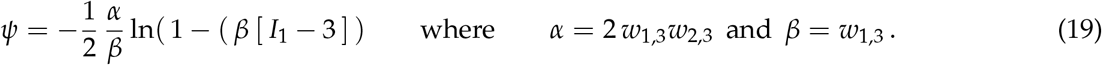

The Gent model translates into the following material table for our universal material subroutine,

> *PARAMETER TABLE, TYPE=“UNIVERSAL_TAB” 1,1,3,w_1,3_,w_2,3_

and activates the third term of our model.

#### Holzapfel model

The Holzapfel model [26] uses quadratic exponentials, typically of the fourth invariant, which we adapt here for the the first invariant, [*I*_1_ − 3], in terms of two parameters *a* and *b*. We recover it as a special case of the network free energy (9) as

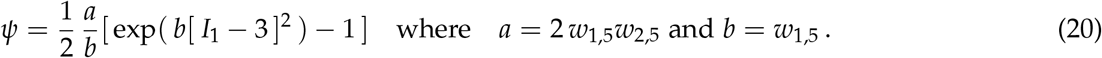

The Holzapfel model translates into the following material table for our universal material subroutine,

> *PARAMETER TABLE, TYPE=“UNIVERSAL_TAB” 1,2,2,w_1,5_,w_2,5_

and activates the fifth term of our model.

Figure 3 compares the neo Hooke, Blatz Ko, Demiray, and Holzapfel models and the finite element simulation with our universal material subroutine. The graphs show the nominal stress as a function of the stretch and shear strain for all four models. The dots indicate the tension, compression, and shear data of human gray matter tissue, the color-coded areas highlights the contributions to the stress function. The finite element simulation with our universal material subroutine in the bottom graphs agrees excellently with the discovered models in the top graphs [35] and confirms the correct implementation of the first, seventh, second, and fifth terms of our model.

**Figure 3:**
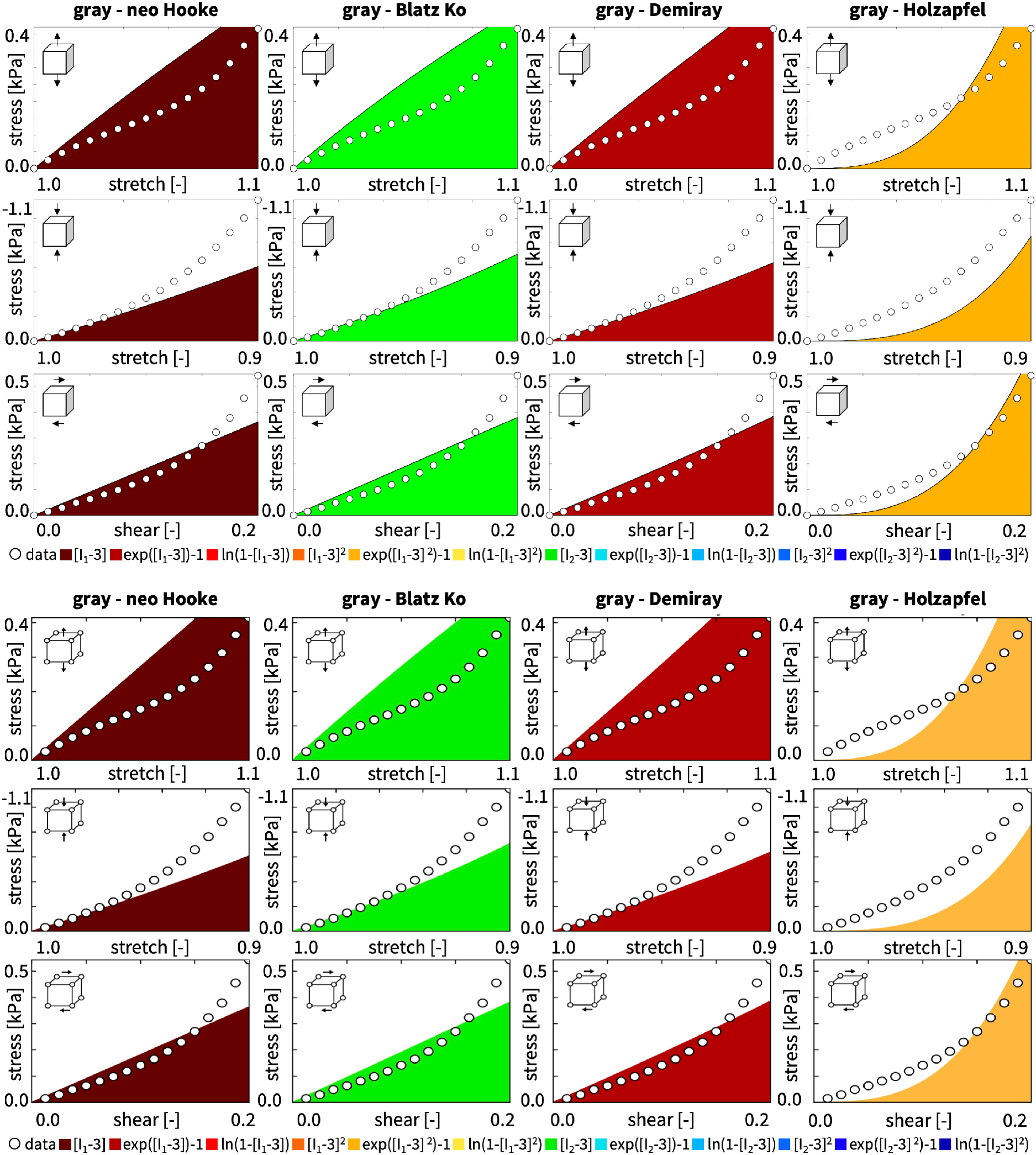
Neo Hooke, Blatz Ko, Mooney Rivlin, Demiray, Gent, and Holzapfel models and finite element simulation. Nominal stress as a function of stretch and shear strain for the neo Hooke model 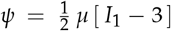 with `1,1,1,w`_1,1_`,w`_2,1_, the Blatz Ko model 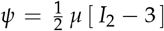 with `2,1,1,w`_1,7_`,w`_2,7_, the Demiray model 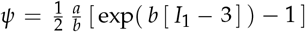 with `1,1,2,w`_1,2_`,w`_2,2_, and the Holzapfel model 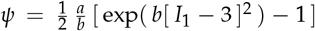 with `1,2,2,w`_1,5_`,w`_2,5_. Dots illustrate the tension, compression, and shear data of human gray matter tissue [35]; color-coded area highlights the contribution to the stress function according to Table 1; top graphs display the discovered model and bottom graphs display the finite element simulation.

### 5.2 Benchmarking with newly discovered models

To illustrate how our universal material subroutine performs for newly discovered models, we benchmark our subroutine with two recently discovered models for gray and white matter tissue [35], translate their network weights *w*_1,•_ and *w*_2, •_ of the first and second layers into their model parameters, provide the material table for the input file to our material subroutine, and compare each simulation to experimental data. Table 2 summarizes our discovered weights *w*_1, •_ and *w*_2, •_ when training our network with individual and combined tension, compression, and shear data from human gray and white matter brain tissue.

**Table 2:** Newly discovered gray and white matter models and parameters. Discovered gray and white matter weights *w*_1,•_ and *w*_2,•_ for training with individual and combined tension, compression, and shear data and for human gray and white matter brain tissue [35].

|  | tension |  | compression |  | shear |  | ten+com+shr |  | tension |  | compression |  | shear |  | ten+com+shr |  |
| --- | --- | --- | --- | --- | --- | --- | --- | --- | --- | --- | --- | --- | --- | --- | --- | --- |
| | gray matter<br>$n = 15$ | | gray matter<br>$n = 17$ | | gray matter<br>$n = 35$ | | gray matter<br>$n = 15, 17, 35$ | | white matter<br>$n = 18$ | | white matter<br>$n = 18$ | | white matter<br>$n = 33$ | | white matter<br>$n = 18, 18, 33$ | |
| | $w_{1,\bullet}$<br>[-] | $w_{2,\bullet}$<br>[kPa] | $w_{1,\bullet}$<br>[-] | $w_{2,\bullet}$<br>[kPa] | $w_{1,\bullet}$<br>[-] | $w_{2,\bullet}$<br>[kPa] | $w_{1,\bullet}$<br>[-] | $w_{2,\bullet}$<br>[kPa] | $w_{1,\bullet}$<br>[-] | $w_{2,\bullet}$<br>[kPa] | $w_{1,\bullet}$<br>[-] | $w_{2,\bullet}$<br>[kPa] | $w_{1,\bullet}$<br>[-] | $w_{2,\bullet}$<br>[kPa] | $w_{1,\bullet}$<br>[-] | $w_{2,\bullet}$<br>[kPa] |
| $w_{\bullet,1}$ | 0.314 | 0.346 | 0.403 | 0.198 | 0.663 | 0.180 | 0.000 | 0.000 | 0.000 | 0.000 | 1.736 | 0.281 | 0.364 | 0.249 | 0.000 | 0.000 |
| $w_{\bullet,2}$ | 0.158 | 0.110 | 0.063 | 0.790 | 0.242 | 0.260 | 0.000 | 0.000 | 0.000 | 0.000 | 0.000 | 0.000 | 0.103 | 0.240 | 0.000 | 0.000 |
| $w_{\bullet,3}$ | 0.000 | 0.000 | 0.000 | 0.000 | 0.766 | 0.184 | 0.000 | 0.000 | 0.000 | 0.000 | 0.000 | 0.000 | 0.037 | 0.307 | 0.000 | 0.000 |
| $w_{\bullet,4}$ | 1.130 | 0.681 | 2.373 | 1.109 | 1.440 | 1.420 | 0.000 | 0.000 | 0.893 | 0.147 | 1.547 | 1.077 | 1.394 | 0.652 | 0.000 | 0.000 |
| $w_{\bullet,5}$ | 1.472 | 1.562 | 1.186 | 2.103 | 1.336 | 1.711 | 0.000 | 0.000 | 0.376 | 0.233 | 1.142 | 1.215 | 1.360 | 1.103 | 0.000 | 0.000 |
| $w_{\bullet,6}$ | 0.502 | 0.435 | 0.000 | 0.000 | 0.000 | 0.000 | 0.000 | 0.000 | 1.308 | 0.430 | 1.212 | 1.148 | 0.440 | 0.831 | 0.000 | 0.000 |
| $w_{\bullet,7}$ | 0.952 | 0.169 | 1.853 | 0.290 | 0.373 | 0.190 | 0.000 | 0.000 | 1.004 | 0.072 | 0.000 | 0.000 | 0.035 | 0.295 | 1.386 | 0.160 |
| $w_{\bullet,8}$ | 0.228 | 0.207 | 0.059 | 0.059 | 0.261 | 0.357 | 0.000 | 0.000 | 0.087 | 0.072 | 0.003 | 0.030 | 0.055 | 0.391 | 0.240 | 0.490 |
| $w_{\bullet,9}$ | 0.682 | 0.173 | 1.947 | 0.114 | 0.000 | 0.000 | 0.988 | 0.634 | 0.840 | 0.207 | 0.000 | 0.000 | 0.768 | 0.118 | 0.000 | 0.000 |
| $w_{\bullet,10}$ | 2.264 | 0.848 | 2.274 | 1.130 | 0.880 | 1.987 | 2.774 | 1.370 | 0.000 | 0.000 | 1.008 | 1.413 | 1.055 | 0.855 | 0.000 | 0.000 |
| $w_{\bullet,11}$ | 0.038 | 0.357 | 1.223 | 2.067 | 1.735 | 1.551 | 1.650 | 1.888 | 0.000 | 0.000 | 1.219 | 1.133 | 0.999 | 1.074 | 1.889 | 1.686 |
| $w_{\bullet,12}$ | 0.933 | 0.473 | 0.000 | 0.000 | 0.882 | 1.425 | 1.403 | 1.666 | 1.105 | 0.003 | 2.648 | 0.823 | 0.000 | 0.000 | 1.179 | 1.911 |

#### Gray matter model

The left columns of Table 2 provide four different models for gray matter tissue, three for training with the individual tension, compression, and shear data, and one for training with all three data sets combined. When trained with the individual data sets for tension, compression, and shear, the neural network in Figure 1 discovers the majority of terms of the free energy function (9), eleven, nine, and ten terms, while only one, three, and two terms train to zero. When trained with all three data sets combined, our network uniquely discovers a four-term model, while the weights of the other eight terms train to zero,

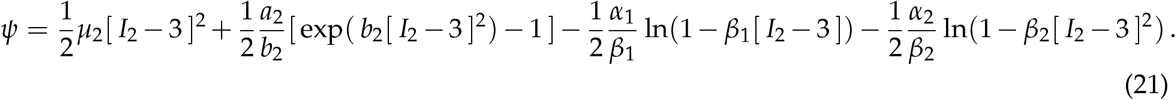

The non-zero weights translate into physically meaningful gray matter parameters with well-defined physical units, the four stiffness-like parameters, *μ*_2_ = 2 *w*_1,10_ *w*_2,10_ = 7.60kPa, *a*_2_ = 2 *w*_1,11_ *w*_2,11_ = 6.23kPa, *α*_1_ = 2 *w*_1,9_ *w*_2,9_ = 1.25kPa, *α*_2_ = 2 *w*_1,12_ *w*_2,12_ = 4.67kPa, and the three nonlinearity parameters, *b*_2_ = *w*_1,11_ = 1.65, *β*_1_ = *w*_1,9_ = 0.99, *β*_2_ = *w*_1,12_ = 1.40. The newly discovered gray matter model translates into the following material table for our universal material subroutine.

> *PARAMETER TABLE, TYPE=“UNIVERSAL_TAB” 2,1,3,w_1,9_,w_2,9_ 2,2,1,w_1,10_,w_2,10_ 2,2,2,w_1,11_,w_2,11_ 2,2,3,w_1,12_,w_2,12_

Figure 4 compares the gray matter model and the finite element simulation with our universal material subroutine. The graphs show the nominal stress as a function of the stretch and shear strain for the gray matter model. The dots indicate the tension, compression, and shear data of human gray matter tissue, the color-coded areas highlight the contributions to the stress function. The finite element simulation with our universal material subroutine in the bottom graphs agrees excellently with the discovered gray matter model in the top graphs [35] and confirms the correct implementation of all twelve terms of our model.

**Figure 4:**
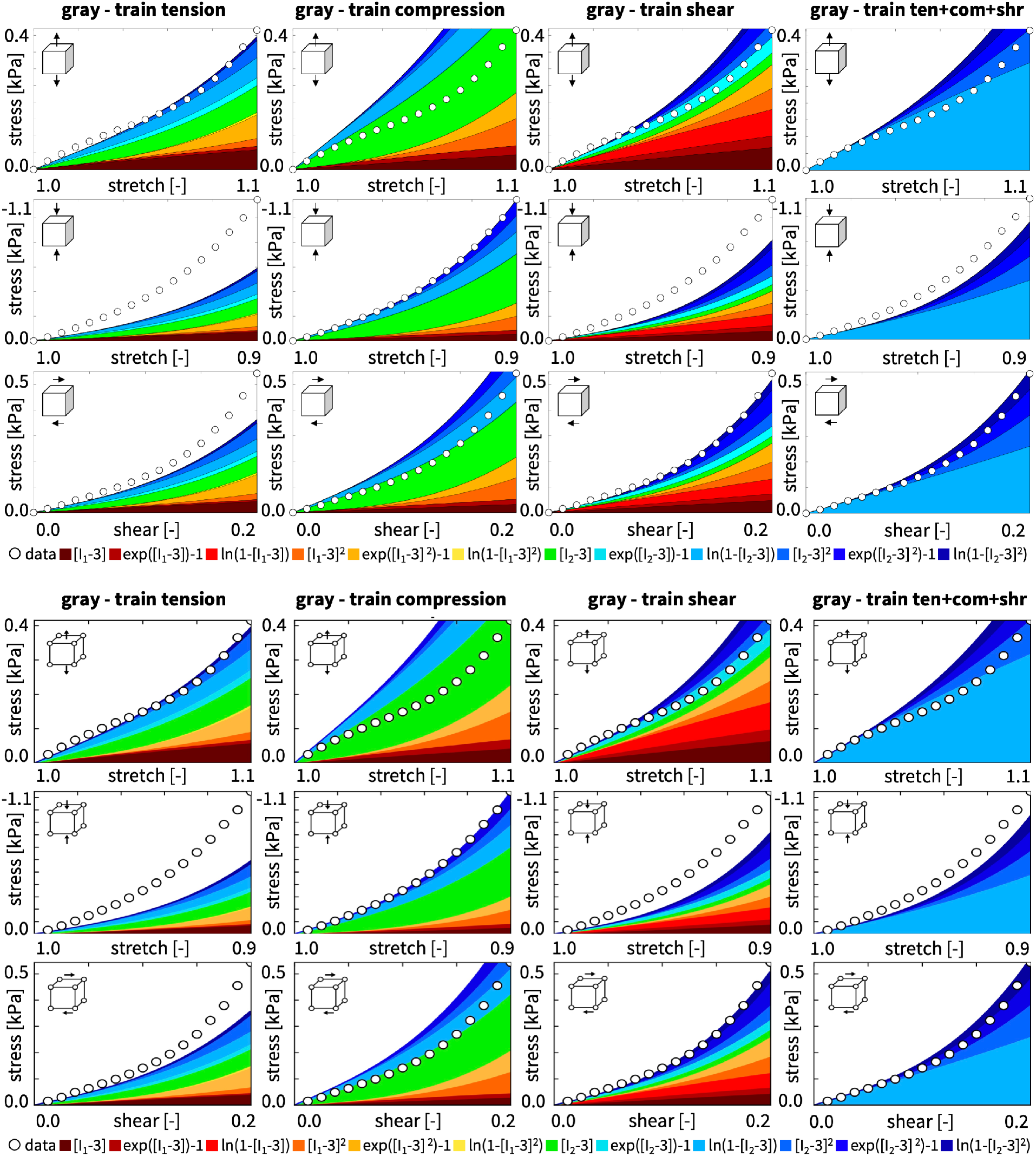
Gray matter discovered model and finite element simulation. Nominal stress as a function of stretch and shear strain for the isotropic, perfectly incompressible constitutive neural network with two hidden layers, and twelve nodes in Figure 1. Dots illustrate the tension, compression, and shear data of gray matter [35]; color-coded areas highlight the twelve contributions to the discovered stress function according to Table 2; top graphs display the discovered model and bottom graphs display the finite element simulation.

#### White matter model

The right columns of Table 2 provide four different models for white matter tissue, three for training with the individual tension, compression, and shear data, and one for training with all three data sets combined. When trained with the individual data sets for tension, compression, and shear, the neural network in Figure 1 discovers the majority of terms of the free energy function (9), seven, eight, and eleven terms, while only five, four, and one terms train to zero. When trained with all three data sets combined, our network uniquely discovers a four-term model, while the weights of the other eight terms train to zero,

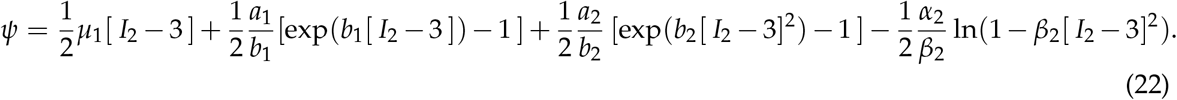

The non-zero weights translate into physically meaningful parameters with well-defined physical units, the four stiffness-like parameters, *μ*_1_ = 2 *w*_1,7_ *w*_2,7_ = 0.44kPa, *a*_1_ = 2 *w*_1,8_ *w*_2,8_ = 0.24kPa, *a*_2_ = 2 *w*_1,11_ *w*_2,11_ = 6.37kPa, *α*_2_ = 2 *w*_1,12_ *w*_2,12_ = 4.51kPa, and the three nonlinearity parameters, *b*_1_ = *w*_1,8_ = 0.24, *b*_2_ = *w*_1,11_ = 1.89, *β*_2_ = *w*_1,12_ = 1.18. The newly discovered white matter model translates into the following material table for our universal material subroutine.

> *PARAMETER TABLE, TYPE=“UNIVERSAL_TAB” 2,1,1,w_1,7_,w_2,7_ 2,1,2,w_1,8_,w_2,8_ 2,2,2,w_1,11_,w_2,11_ 2,2,3,w_1,12_,w_2,12_

Figure 5 compares the white matter model and the finite element simulation with our universal material subroutine. The graphs show the nominal stress as a function of the stretch and shear strain for the white matter model. The dots indicate the tension, compression, and shear data of human white matter tissue, the color-coded areas highlight the contributions to the stress function. The finite element simulation with our universal material subroutine in the bottom graphs agrees excellently with the discovered white matter model in the top graphs [35] and confirms the correct implementation of the twelve terms of our model.

**Figure 5:**
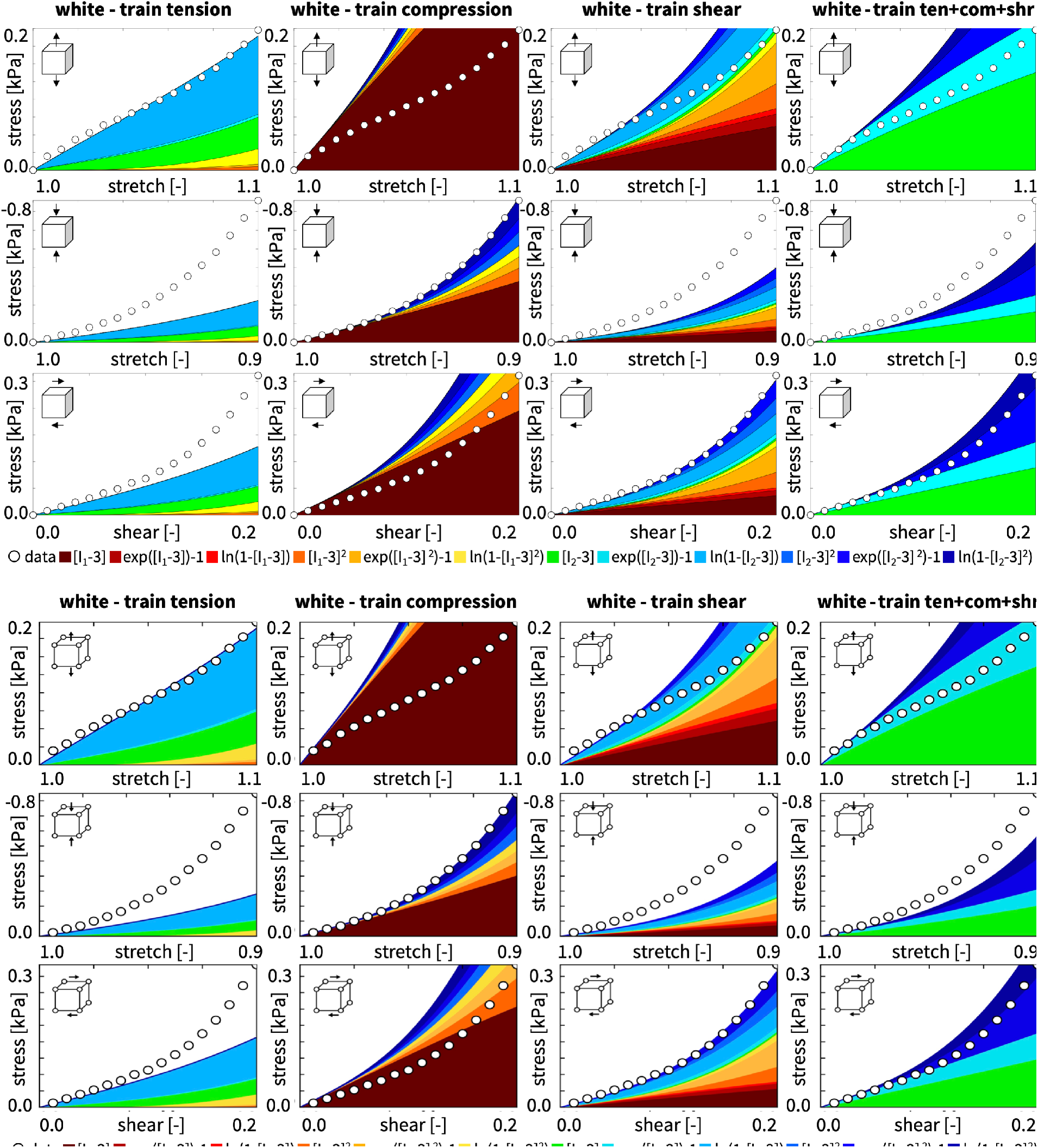
White matter discovered model and finite element simulation. Nominal stress as a function of stretch and shear strain for the isotropic, perfectly incompressible constitutive neural network with two hidden layers, and twelve nodes in Figure 1. Dots illustrate the tension, compression, and shear data of white matter [35]; color-coded areas highlight the twelve contributions to the discovered stress function according to Table 2; top graphs display the discovered model and bottom graphs display the finite element simulation.

Taken together, these eight examples demonstrate that our proposed method generalizes well to previously undiscovered constitutive functions, which translate smoothly into a universal material subroutine that agrees well with the experimental data and previous simulations.

### 5.3 Realistic finite element simulations

To illustrate the performance of our universal material subroutine for our discovered gray and white matter models from equations (21) and (22) within a realistic finite element simulation, we study six different head impact scenarios. Figure 6 shows our sagittal model that consists of 6182 gray and 5701 white matter linear triangular elements, 6441 nodes, and 12,882 degrees of freedom. Figure 7 shows coronal model that consists of 7106 gray and 14196 white matter linear triangular elements, 11808 nodes, and 23616 degrees of freedom. We embed both models into the skull using spring support at the free boundaries and apply top-of-the-head, diagonal, and frontal impacts to the sagittal model and top-of-the-head, diagonal, and lateral impacts to the coronal model.

**Figure 6:**
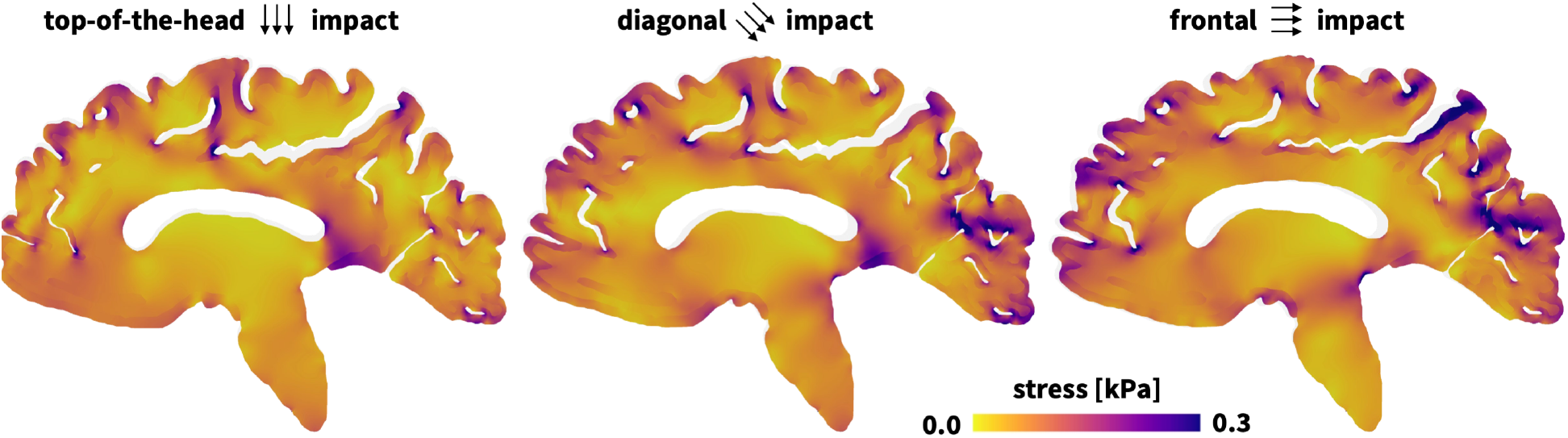
Stress profiles for top-of-the-head, diagonal, and frontal impact to the human brain. The finite element simulations use our universal material subroutine with the discovered models from Table 1 for gray matter from equation (21) with the four stiffness-like parameters *μ* = 7.60kPa, *a*_2_ = 6.23kPa, *α*_1_ = 1.25kPa, *α*_2_ = 4.67kPa, and the three nonlinearity parameters, *b*_2_ = 1.65, *β*_1_ = 0.99, *β*_2_ = 1.40, and for white matter from equation (22) with the three stiffness-like parameters *μ* = 0.44kPa, *a*_1_ = 0.24kPa, *a*_2_ = 6.37kPa, *α*_2_ = 4.51kPa, and the three nonlinearity parameters, *b*_1_ = 0.24, *b*_2_ = 1.89, *β*_2_ = 1.18.

**Figure 7:**
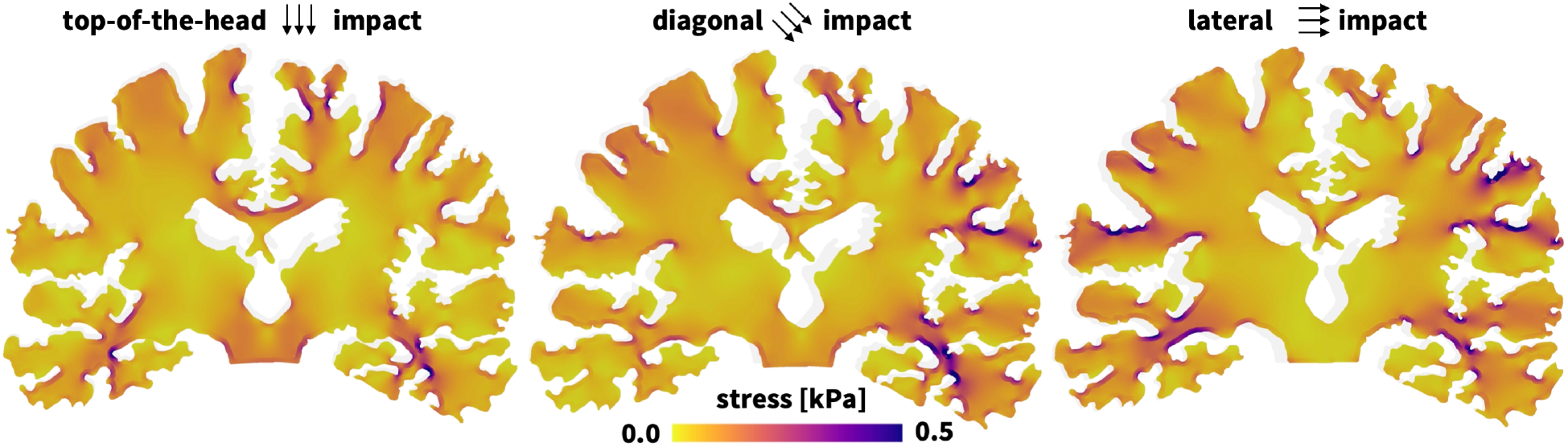
Stress profiles for to top-of-the-head, diagonal, and lateral impact to the human brain. The finite element simulations use our universal material subroutine with the discovered models from Table 1 for gray matter from equation (21) with the four stiffness-like parameters *μ* = 7.60kPa, *a*_2_ = 6.23kPa, *α*_1_ = 1.25kPa, *α*_2_ = 4.67kPa, and the three nonlinearity parameters, *b*_2_ = 1.65, *β*_1_ = 0.99, *β*_2_ = 1.40, and for white matter from equation (22) with the three stiffness-like parameters *μ* = 0.44kPa, *a*_1_ = 0.24kPa, *a*_2_ = 6.37kPa, *α*_2_ = 4.51kPa, and the three nonlinearity parameters, *b*_1_ = 0.24, *b*_2_ = 1.89, *β*_2_ = 1.18.

Figures 6 and 7 summarize the stress profiles for the six different impact simulations. Clearly, we observe stress concentrations at the gray and white matter interface, between the cortex and the corona radiata. These stress concentrations are common after a hit to the head, and are a result of the structural and mechanical differences between different tissue types: Gray matter consists primarily of neuronal cell bodies and is rather dense, while white matter consists primarily of myelinated axons. Upon an impact to the head, forces are transmitted differently through these tissue types. From Figures 4 and 5, we conclude that gray matter is almost twice as stiff as white matter, with maximum tensile stresses of 0.4 kPa versus 0.2 kPa for stretches of 1.1, maximum compressive stresses of -1.1 kPa versus -0.8 kPa for stretches of 0.9, and maximum shear stresses of 0.5 kPa versus 0.5 kPa for shear of 0.2. This disparity in mechanical stiffnesses leads to localized stress concentrations at the gray and white matter interface, which can disrupt the structural integrity of the tissue and trigger diffuse axonal injuries. The simulations predict that these stress concentrations occur primarily in the frontal and occipital lobes for top-of-the-head impacts, in the deep white matter tracts for diagonal impacts, in the frontal and parietal lobes for frontal impacts, and in the gray and white matter interface for lateral impacts.

Taken together, these six examples demonstrate that our discovered gray and white matter models translate smoothly into a universal material subroutine that generalizes from the homogeneous simulations in Figures 3 through 5 to realistic finite element simulations in Figures 6 and 7, where it robustly predicts heterogeneous stress profiles across complex structures.

## 6 Discussion

### Our universal material subroutine specializes well to popular constitutive models

To demonstrate that our material subroutine includes popular constitutive models as special cases, we benchmarked it with four widely used models. Figure 3 compares the neo Hooke [60], Blatz Ko [6], Demiray [12], and Holzapfel [26] models in the top row to finite element simulations with our universal material subroutine in the bottom row. All four models only activate a single term of the subroutine, which translates into a single-row material table, and a single-color stress plot. For all four models, the finite element simulation with our new material subroutine in the bottom row agrees excellently with the model in the top row [35]. These simple benchmark examples demonstrate that we can recover popular constitutive models for which the weights of our constitutive neural network in Figure 1 gain a well-defined physical meaning and the universal material subroutine takes the functional form of one of the twelve activation functions in Figure 2 [34]. Importantly, to perform a finite element analysis, we no longer need to select a specific material model; instead, we can simply use our universal material subroutine and selectively activate its relevant terms through the non-zero entries in the material table.

### Our universal material subroutine expands naturally to compressible and anisotropic materials

For illustrative purposes, we have only demonstrated the versatility of our material subroutine for *incompressible* and *isotropic* hyperelastic materials [34]. For these, the first index of our parameter table selects between the first and second invariants, the second index raises them to linear or quadratic powers, and the third index selects between the identity, exponential, and logarithmic functions. This setting seamlessly generalizes to *compressible* and *anisotropic* materials by selecting a first index of three, four, five, …, `NINV` to include terms in the third, fourth, or fifth invariants. For example, the subroutine `UANISOHYPER_INV` supports up to three fiber directions resulting in a total of 15 invariants. It also naturally allows for higher order powers by selecting a second index larger than two and facilitates the integration of additional functional forms through a third index larger than three. In the present study, we have illustrated how to translate the output of our automated model discovery into the input of our universal material subroutine within Abaqus [1] using the software’s invariant-based user material subroutine `UANISOHYPER_INV`. However, the inherent modularity of our approach ensures that this translation will generalize naturally to arbitrary implicit or explicit nonlinear finite element packages.

### Our proposed method generalizes well to previously undiscovered constitutive functions

Automated model discovery allows us to discover the best possible model, in our case out of 2^12^ = 4096 possible combinations of terms [35]. Traditionally, model developers have rationalized constitutive models from the shape of experimental curves and then fit their parameters to data. Throughout the past decades, this has generated dozens of models with one [6, 12, 19, 60], two [26, 42, 49], three [64] or more terms, almost always in terms of the *first invariant*. Recent developments in deep learning now allow us to rapidly screen thousands of possible combinations of terms and discover the best possible fit. However, when only trained with individual tension, compression, or shear data, the network tends to overfit the data and discovers a wide variety of terms [34, 54]. Yet, when trained with all three data sets combined, the network robustly and repeatedly discovers a small subset of terms in the *second invariant* for both gray and white matter [35]. Strikingly, these terms have been overlooked by traditional manual model development. In retrospect, it seems obvious that the second invariant is well suited to characterize human brain tissue : While the first invariant, 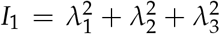, is quadratic in terms of the stretches *λ*, the second invariant, 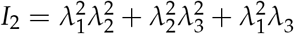, is quartic and seems better suited to represent nonlinearities [29]. This is particularly relevant in the small stretch regime of 0.9 ≤ *λ* ≤ 1.1 that we study here, where the stretches are small and their nonlinear effects remain minor [10]. For rubber-like materials, where the stretches in uniaxial tension, equibiaxial tension, and pure shear can easily reach values of 1.0 ≤ *λ* ≤ 8.0, the second invariant explodes and seems less well-suited to characterize the stretch-stress response [59]. The excellent agreement of the finite element simulations with our universal material subroutine in the bottom rows of Figures 4 and 5 with the experimental data [10] and the discovered models [35] in the top rows confirms the correct implementation of our discovered gray and white matter models.

### Our universal material subroutine uses interpretable material parameters

A characteristic feature of our proposed modeling strategy is that it features different activation functions, linear and quadratic, (◦)^1^ and (◦)^2^, embedded in the identity, exponential, and logarithmic functions, (◦), (exp(◦) −1), and (−ln(1 − (◦))) [34]. This is in stark contrast to previous approaches that have used one and the same activation function across all network nodes, for example of hyperbolic tangent [24, 56], exponential linear unit [31], or softplus squared [4] type. While it is theoretically possible to manually embed these models into a finite element workflow, their weights translate into material parameters that have no clear physical interpretation [32]. In contrast, our non-zero weights translate into physically meaningful parameters with well-defined physical units: the stress-like parameters, *w*_1,•_ *w*_2,•_, with the unit kilopascal, and the dimensionless parameters, *w*_1,•_, that govern the exponential [12] and logarithmic [19] nonlinearities. We conclude that our proposed approach generalizes well to previously undiscovered constitutive functions, which translate naturally into a universal material subroutine that agrees well with the experimental data [10] and with previous simulations [34]. Importantly, rather than having to implement a new material subroutine for each newly discovered model, we use a single universal material subroutine that inherently incorporates all 2^12^ = 4096 possible combinations of terms and activates the relevant model merely by means of the twelve rows of its parameter table.

### Our universal material subroutine generalizes well to realistic simulations

When embedded into a finite element simulation, our material subroutine translates the local deformation gradient into stresses and stress derivatives that enter the global force vector and stiffness matrix of the local Newton iteration to solve the balance of motion. To illustrate that our new subroutine not only performs well for the homogeneous examples in Figures 3 to 5, but also for realistic finite element simulations, we simulate the regional stress distributions across the human brain for six different head impact scenarios [44]. Figures 6 and 7 emphasize the sensitivity of the stress profiles with respect to the location and direction of the impact. Depending on impact location and severity, individuals may experience a broad spectrum of symptoms ranging from headaches, dizziness, nausea, and vision problems to difficulties with concentration and attention [22]. *Top-of-the-head impacts* in Figures 6 and 7, left, affect regions of the skull that are usually very thin and vulnerable to skull fractures, brain contusions, and significant brain damage [52]. In agreement with our simulated stress profiles, these impacts primarily affect the frontal region of the brain that plays a crucial role in higher cognitive functions, personality, emotional regulation, and decision-making. Their symptoms may range from cognitive impairment, memory loss, and motor function deficit to long-term consequences such as permanent disability or death. *Diagonal impacts* as in Figures 6 and 7, middle, can cause rotational forces that many result in diffuse axonal injuries. These injuries occur when brain structures tear in response to elevated shear stresses [20]. Diffuse axonal injuries often involve deep white matter tracts and can affect multiple lobes of the brain, including the frontal, temporal, and parietal lobes, for which our simulation predicts elevated stress levels. These can have profound effects on brain function and lead to cognitive, behavioral, and motor impairments associated with difficulties of attention, memory, problem-solving, and emotional regulation. *Frontal impacts* as in Figures 6, right, directly affect the frontal region of the brain, the side of impact, through coup injury. Importantly, they can also have severe secondary effects on brain regions opposite to the impact, through countercoup injury, as we conclude from our simulated stress profiles. Frontal impacts commonly results in mild or severe concussions [22] or traumatic brain injuries associated with a wide range of symptoms such as headaches, dizziness, confusion, and memory issues. These can impair higher cognitive functions, personality, and emotional regulation and lead to changes in behavior, mood, and decision-making. *Lateral impacts* as in Figures 7, right, cause the brain to rotate, which can induce diffuse axonal injuries similar to diagonal impacts [9]. As we conclude from our simulated stress profiles, lateral impacts affect mainly the gray and white matter interface, which experiences much higher stresses than, for example, under top-of-the-head impacts of the same magnitude. Diffuse axonal injuries can result in cognitive, behavioral, and motor impairments, and affect various aspects of daily life. Lateral impacts may also lead to contusions, which can compromise brain function and potentially cause long-term memory loss, personality changes, and emotional instability. Knowing the precise location and direction of a head impact is critical because impacts to different brain regions can result in varying types and severity of injuries [47]. Understanding the stress profiles in response to different types of impact can help assess the extent of an injury, determine the appropriate treatment, and develop strategies to prevent further head trauma [63].

## Limitations

Our results suggest that we can seamlessly integrate automated model discovery into a finite element workflow through a new universal material subroutine. Nonetheless, our study has several limitations that point towards possible future extensions. First, while our current model is incompressible and isotropic, we can easily expand it to include compressibility [21] and anisotropy [51] by adding the third, fourth, fifth, and higher order invariants, that we can simply embed via the first index in our parameter table. Second, we can expand our model and include higher order powers [64], cubic or quartic, via the second index in our parameter table. Third, we could generalize our current network architecture from an additive coupling of the invariants towards a multiplicative coupling [18], which would translate into additional cross-coupling terms in the tangents of our user material subroutine. Fourth, instead of using a purely invariant-based formulation, we could also include principal-stretch-based terms [54] that mimic an Ogden [45] or Valanis-Landel [61] type behavior. Fifth, in addition to the elastic potential that characterizes the hyperelastic behavior, we could also include one or more inelastic potentials that characterize viscosity, plasticity, damage, or growth [62].

## 7 Conclusion

Constitutive modeling is critical to a successful analysis of materials and structures. However, the scientific criteria for selecting the appropriate model remain insufficiently understood. This work seeks to address the question whether and how we can automate constitutive modeling within a finite element analysis. Our work is made possible by a recent trend in physics-based artificial intelligence, automated model discovery, a new technology that allows us to autonomously discover the best model to explain experimental data. Automated model discovery comes in various flavors and uses sparse regression, genetic programming, or constitutive neural networks with the common goal to discover constitutive models from thousands of combinations of a few functional building blocks. Here our objective was to integrate automated model discovery into the finite element workflow by creating a single unified user material subroutine that contains 2^12^ = 4096 constitutive models made up of 12 individual terms. For illustrative purposes, we prototyped this strategy within the `UANISOHYPER_INV` environment of the general-purpose finite element software Abaqus and share our new universal material subroutine publicly on GitHub. For three examples, we demonstrated that our universal material subroutine specializes well to traditional constitutive models, generalizes well to newly discovered models, and performs well within realistic finite element simulations. While we have only prototyped our approach for a specific hyperelastic material model, for a specific type of automated model discovery, and for a specific finite element platform, we are confident that our strategy will generalize naturally to more complex anisotropic, compressible, and inelastic materials, to other types of model discovery, and to other nonlinear finite element analysis platforms. Replacing dozens of individual material subroutines by a single universal material subroutine–populated directly via automated model discovery–makes finite element analyses more accessible, more robust, and less vulnerable to human error. This could induce a paradigm shift in constitutive modeling and forever change how we simulate materials and structures.

## Data availability

Our neural network, data, and universal material subroutine will be available upon request or at https://github.com/LivingMatterLab/CANN.

## Acknowledgments

This work was supported by a DAAD Fellowship to Kevin Linka, and by NSF CMMI Award 2320933 *Automated Model Discovery for Soft Matter* to Ellen Kuhl.

## Notes

### Competing Interest Statement

The authors have declared no competing interest.

